# Inactive X chromosome DNA methylation erosion with age in human dermal fibroblasts

**DOI:** 10.64898/2026.11.19.767824

**Authors:** Jannat Ijaz, Ioannis Kafetzopoulos, Naomi Park, Felix Krueger, James Clarke, Jill Temple, Renato Colaco, Marta Fernández de Ara, Paul Coupland, Ricard Argelaguet, Aled Parry, Ariane Lismer, Jasmin Taubenschmid-Stowers, Sergei A. Manakov, Fatima Santos, Wolf Reik

## Abstract

X chromosome inactivation is an epigenetic process that establishes dosage compensation between the sexes through the monoallelic silencing of one X chromosome in karyotypically normal females. The allele that becomes inactivated is determined randomly during early development, resulting in females being mosaic with respect to whether the maternal or paternal allele is inactivated. This mosaicism makes it technically challenging to separate the active and inactive chromosomes in bulk populations, and consequently it has been historically difficult to study X chromosome inactivation in human cells. Here we use long read sequencing of 31 single cell clones derived from primary human dermal fibroblasts from 7 donors aged 25-80 years, and generate a method to distinguish DNA methylation state of the active versus the inactive X chromosome. As expected, CpG islands (CGIs) on the inactive X chromosome have high levels of methylation, while CGIs on the active X chromosome are very lowly methylated. Notably, we observe substantial loss of DNA methylation across multiple CGIs on the inactive X chromosome in an age dependent manner. We also observe this DNA methylation erosion in bulk samples by quantifying CGI methylation levels across individual sequencing reads. The genes that show age-related erosion of CGI methylation are associated with increased total transcription suggesting this methylation erosion generates a transcriptional permissive state. Overall, our study provides evidence of erosion of X chromosome inactivation with age in cell lines derived from primary human samples and implicates a role for loss of DNA methylation in the increased dosage of X chromosome linked genes in females with age. These findings shed new light on the epigenetic instability of the inactive X chromosome during human ageing and may be linked to age-related cellular dysfunction in females.

## Introduction

The difference in size between the X and Y chromosomes and the low number of genes located on the Y chromosome mean that biological women (46, XX) carry roughly double the genetic information to biological men (46, XY) on their sex chromosomes. This difference necessitates a dosage compensation mechanism where one X chromosome gets extensively repressed in women by a process called X chromosome inactivation (XCI). XCI was first proposed in 1961 (Lyon 1961) and is an essential mechanism for dosage compensation in placental mammals (Dossin and Heard 2022; Alfeghaly and Rougeulle 2025). In humans, XCI occurs during early development and is random with respect to the parental alleles, meaning that tissues have a mix of maternal and paternal alleles inactivated (Amos-Landgraf et al. 2006; Novelli et al. 2003; Gylemo et al. 2025). XCI occurs on the chromosome where the long non-coding RNA XIST is upregulated and leads to the establishment of repressive histone modifications and DNA methylation (Zhao et al. 2008; Chu et al. 2015). In particular, CpG islands across the inactive X chromosome are highly methylated, particularly by DNMT3B, and transcriptionally silenced, which has been hypothesised to serve as a permanent epigenetic ‘lock’. Following the random establishment of the inactive allele, the pattern is stable and heritable through subsequent cell divisions (Lock et al. 1987; Hansen et al. 1999; Sharp et al. 2011; Gendrel et al. 2012)

The process of inactivation does not repress all genes on the X chromosome. There are constitutive escapee genes which are never repressed and facultative escapees which have variable repression (Tukiainen et al. 2017). Constitutive escapees include the pseudoautosomal region that is present on the X and the Y chromosomes so must evade repression to achieve the same expression levels in both sexes. Other constitutive escapees are not on the Y chromosome so their increased dosage may contribute to differences in phenotypes between the sexes, including disease susceptibility, onset and progression. Facultative escapees vary in different tissues and have been shown to escape in a cell, tissue or individual specific manner. This has been shown in humans using both bulk RNA-seq (Tukiainen et al. 2017), and single cell RNA-seq (Gylemo et al. 2025). Furthermore, reports of age-related escapees in mice suggest a gradual loss of dosage compensation that may contribute to differences in disease susceptibility, onset and progression between the sexes for diseases that increase with age (Hoelzl et al. 2025; Gadek et al. 2025).

The first evidence of age-related escapees was reported in 1987 when the re-expression of a single gene was seen in the liver of mice with non-random X inactivation (Wareham et al. 1987). Since this initial study, age-related escape from X chromosome inactivation has been the focus of a number of studies in mice using both phenotypic and molecular readouts. Historically it has been easier to study the X chromosome in mice as genetic manipulation methods can be used to force a specific parental X chromosome to become inactivated. Typically this is done by deletion of the *Xist* promoter in one of the two parental chromosomes, meaning that chromosome is never repressed (Hoelzl et al. 2025; Gadek et al. 2025). This allows conventional phasing techniques to be applied to sequencing data to study how gene expression, chromatin accessibility and DNA methylation changes on the inactive X chromosome in different tissues and with age. Observations have included a change in coat colour of mice (S. Brown and Rastan 1988); changes in localisation and expression of *Xist* in mouse hematopoietic stem cells (HSCs) (Grigoryan et al. 2021), increased X chromosome gene expression variability, hypomethylation and increased chromatin accessibility in mouse HSCs (Grigoryan et al. 2021), and age-related escapees in heart, lung, liver, brain, spleen, kidney and muscle in allele-specific Xist knock out mice (Gadek et al. 2025; Hoelzl et al. 2025).

Studying XCI in humans has largely relied on RNA sequencing to look at the allelic ratio of the active vs inactive chromosomes within the same cell. Both bulk (Rocca et al. 2026) and single cell methods (Grigoryan et al. 2021) have been used but they are typically based on short-read technologies meaning differentiating the alleles is challenging and gives only a fragmented view of the transcriptome. Single cell approaches have the benefit of increased power, as thousands of cells can be studied in a single experiment. However, single-cell data is subject to gene dropouts, which make it considerably noisier than bulk measurements. Bulk RNA sequencing, on the other hand, captures more genes but may be underpowered to pick up smaller changes in allelic ratio. Other studies have used methylation array technologies to identify methylation changes with age on the X chromosome (Liu et al. 2023; Kananen and Marttila 2021; Li et al. 2020). However, these studies lack the resolution to identify whether the changes are happening on the active or the inactive chromosomes. There are no studies to date that have the resolution to identify age-related differences in the active compared to the inactive X chromosome methylome in humans.

In this study, we isolated and expanded single cell derived clonal populations from primary human dermal fibroblast cell lines from donors aged 25-80 years old. This naturally aged model allows us to study changes in XCI with age using bulk sequencing as every cell in the clonal population will have the same parental allele inactivated. We performed long-read sequencing with 5mC basecalling of these clones and the bulk cell lines they were derived from and investigated changes on both the active and inactive X chromosomes. We used these fibroblast populations to identify whether there are systematic changes of DNA methylation on the X chromosome with age.

## Results

### Separating the active and inactive X chromosome reads

Heterozygous SNPs can be used to separate maternal and paternal haplotypes (Figure 1A). In non-clonal populations, human dermal fibroblasts are heterogeneous with respect to which parental X chromosome allele is inactivated. Therefore when segregating reads using genotype, the two alleles show very similar methylation profiles (Figure 1B). To ensure the same allele is inactivated across the population, we utilised single-cell derived clonal expansion. Cell lines can be immortalised to derive single cell clonal populations, however this immortalisation can lead to disrupted epigenetic regulation that may mask age-related changes. To preserve the integrity of the ageing epigenome, we supplemented our culture media with fibroblast growth factor 2. This successfully bypassed early-passage replicative senescence and enabled single cell clone expansion to sufficient cell numbers for deep molecular profiling, without the confounding effects of viral or chemical immortalisation. We derived clonal populations from 7 human dermal fibroblast cell lines isolated from donors of different ages, generating 3 to 5 clones per line (31 total clones). We performed long-read genome and transcriptome sequencing of these clones using nanopore sequencing. Variant allele fractions and methylation profiles in these samples were characteristic of clonal populations consistent with high somatic stability of methylation patterns (Supplementary Figure 1A, B). Since the active or repressed state of a specific X chromosome is stably propagated upon cell division, this resulted in the same parental X chromosome allele being inactivated within each clone. Given this, reads can be assigned to the parental allele that they were derived from using heterozygous variants in a process known as phasing and the active and inactive methylation patterns can be investigated.

**Figure 1.**
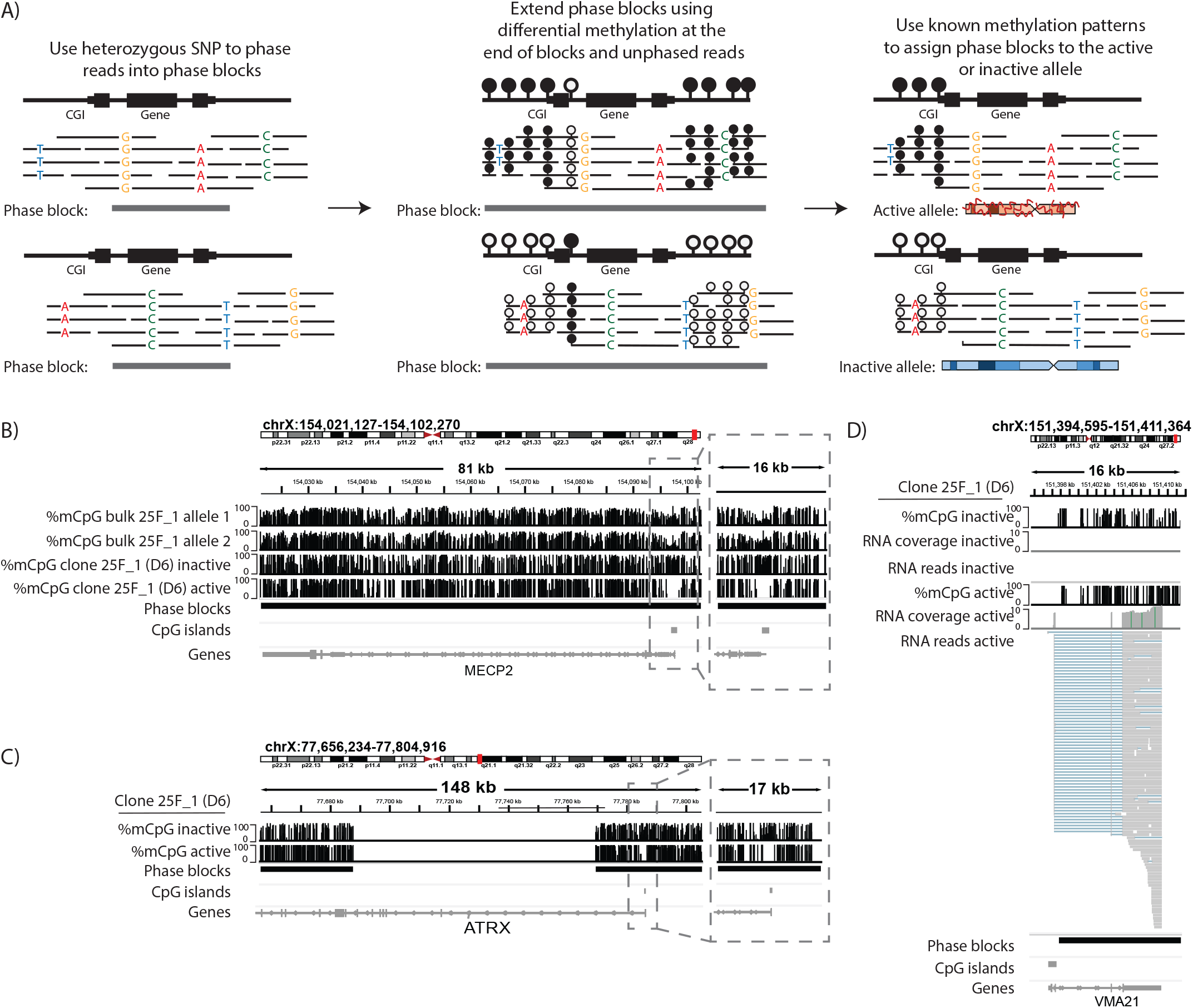
A) Haplotype phasing method schematic. Reads are phased using heterozygous SNPs. Phase blocks were extended using methylation differences at the edge of phase blocks. Phase blocks were assigned to the active or inactive allele using known differences in methylation patterns on the X chromosome. B) Genome browser view of the chrX:154,021,127-154,102,270 including MECP2 gene, an example region that is phased by CpG island differences. CpG methylation from nanopore sequencing is shown with tracks bounded at 0% and 100% methylation. First and second track are the non-clonal lines phased using heterozygous SNPs, with intermediate methylation at the CPG island. Third and fourth tracks are the clonal lines phased using heterozygous SNPs, with distinct methylation patterns at the CPG island. The fifth track is the phase block, showing the region has been phased into a single block. The sixth track is the CpG island annotation and the seventh track is RefSeq gene annotation (also shown as the last three tracks for C and D). The region around the CpG island of the gene has been expanded showing chrX:154,090,341-154,106,585. C) Genome browser view of the chrX:77,656,234-77,804,916 including ATRX gene, an example region that is phased using gene body assignment. CpG methylation from nanopore sequencing is shown with tracks bounded at 0% and 100% methylation. First and second tracks are the inactive and active tracks, respectively. The region around the CpG island of the gene has been expanded showing chrX:77,776,942-77,794,340. D) Genome browser view of the chrX:151,394,595-151,411,364 including VMA21 gene, an example region that is phased using transcription. CpG methylation from nanopore sequencing is shown with tracks bounded at 0% and 100% methylation. The first and fourth second track is the methylation of the inactive and active allele, respectively. The second and fifth track show RNA coverage of the inactive and active allele, respectively. The third and sixth track show raw reads at this locus. Expression is only seen on the active allele.

To identify DNA methylation and transcription patterns on the X chromosome in these clones, we phased sequence variants in nanopore reads using whatshap (Patterson et al. 2015). This led to 71.2% of the X chromosome being phased (standard deviation ± 4.0%) (Figure 1A, Supplementary Figure 1C). We further extended the phase blocks using methphaser (Fu et al. 2024) in regions without sequence variants but with distinct methylation patterns at the edges of the phase blocks. This increased the total amount of the X chromosome that was phased to 79.4% (standard deviation ± 4.3%) (Figure 1A, Supplementary Figure 1C). From this, it is not possible to know how adjacent phase blocks are related to one another. Therefore we used knowledge of expected differences in methylation profiles between the active and inactive allele to group phase blocks, and the reads assigned to them, together (see methods). Specifically, methylation patterns in CpG islands (CGIs) show the most extreme difference between the alleles, making them highly informative for assignment. When phase blocks contained CGIs that were lowly methylated on one allele and highly methylated on the other, we assigned the allele with the high methylation to the inactive allele (Figure 1B). This accounted for 90.0% of the total number of phase blocks we were able to assign (standard deviation ± 5.03%, Supplementary Figure 1D). Some phase blocks did not span CGIs but did share a gene with the adjacent phase block which was assigned. Therefore, we used the higher methylation levels over the gene bodies of actively transcribed genes (Ball et al. 2009) as a pattern to differentiate between the two alleles and link to adjacent phase blocks. If the methylation pattern over the gene body was different between the alleles but similar across the phase blocks, the phase block without a CGI was assigned based on the phasing of the adjacent phase block (Figure 1C). This accounted for 7.84% of the total number of phase blocks we were able to assign (standard deviation of ± 4.57%, Supplementary Figure 1D). Furthermore if there were no informative methylation features in a phase block, SNPs were called in transcripts and genes which contained SNPs were used to assign to phase blocks. If genes were expressed from only one allele, the allele with expressed genes was assigned as active (Figure 1D). This accounted for very few phase blocks being assigned (2.38% with a standard deviation ±1.53%, Supplementary Figure 1D). Some phase blocks, typically in gene-poor regions of the chromosome, did not contain informative features to confidently assign so these were excluded from downstream analysis. Collectively, this led to 40.5% of the X chromosome being confidently assigned to the active or inactive chromosome, respectively (standard deviation ± 7.1%) (Supplementary Figure 1C, 1E). Proportion of assigned bases showed no relationship with donor age (Supplementary Figure 1F, Pearson r = –0.11, p-value = 0.56), indicating that variation in phasing completeness would not confound the age associated methylation changes described below. As expected, the number of bases assigned positively correlated with the input read length (Supplementary Figure 1G, Pearson r = 0.49, p-value=7.3×10^-3^). Phaseable regions of the X chromosome were strongly enriched for genes, GC content, CpG density, and CpG islands relative to non-phaseable regions (Supplementary Figure 1H), consistent with phasing and allele assignment relying on CGI and gene body methylation.

Our method produced methylation patterns across the alleles that were broadly consistent with expected patterns (Sharp et al. 2011). To quantify this, we calculated the average methylation over each phaseable CGI on either allele (mean number of CGIs per clone was 359 with a standard deviation ± 25, maximum possible number was 500). Most of the CGIs were very different between the alleles (Supplementary Figure 2A), with the inactive allele being highly methylated and the active allele being lowly methylated, as expected (Supplementary Figure 2B,C). A subset of CGIs had similar methylation profiles. This was either due to methylation on both alleles as they regulate genes that are repressed on the active allele (2.8% of CGIs per clone, standard deviation ± 0.71%) or due to constitutive escape from repression (6.8% of CGIs per clone, standard deviation ± 1.8%). Interestingly, the promoter associated with *Xist* was the only region that showed low methylation on the inactive allele and high methylation on the active allele, consistent with its expression mediating repression (Supplementary Figure 2A) (Sharp et al. 2011; Trinh et al. 2025). We also observed expected expression patterns as the inactive allele had fewer total counts when compared to the active allele (paired t-test: t-statistic=11.33, p-value=2.3×10^-12^, Supplementary Figure 2D).

### DNA methylation erosion with age in clonal populations

We next compared one clone derived from our youngest donor (25 years old) and one clone derived from our oldest donor (80 years old) to investigate age-related changes in methylation patterns. When looking at the active allele, most CGIs in both the clone derived from the young donor and the clone derived from the old donor were lowly methylated, without exhibiting any substantial age-related changes (Figure 2A, Pearson r = 0.9, p-value<2.2×10^-16^). Strikingly, when looking at the inactive allele, many CGIs that are highly methylated in the clone derived from the young donor exhibited loss of methylation in the clone derived from the old donor (Figure 2B, Pearson r = 0.7, p-value<2.2×10^-16^). This methylation erosion occurred throughout the CGI (example gene in Figure 2C), suggesting that some genes on the inactive X chromosome lost their repressive methylation with age.

**Figure 2.**
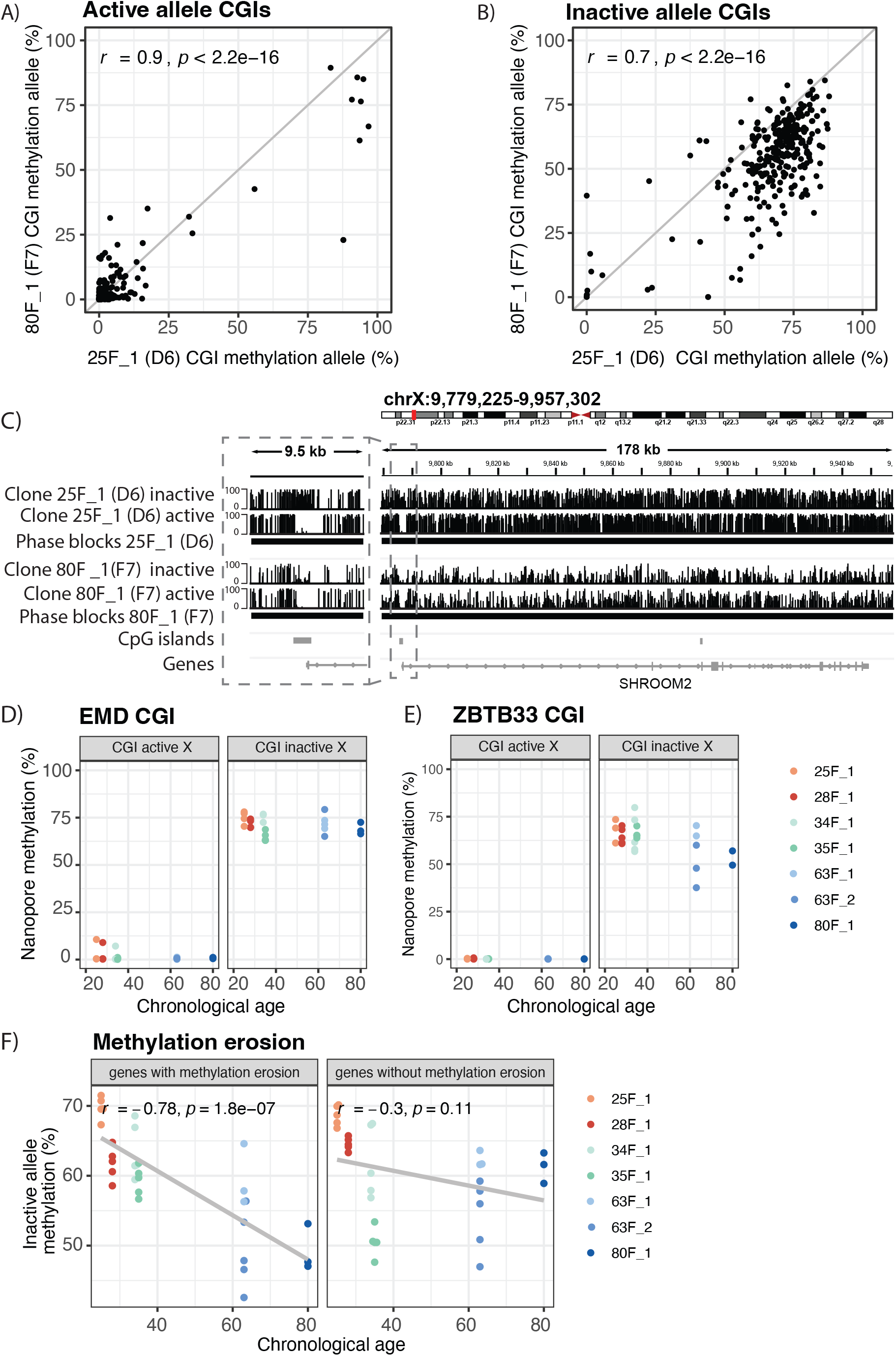
A-B) Mean methylation over all phaseable CpG islands on the active and inactive allele, respectively. Each point is the average methylation over the entire CGI annotation. x=y line shown in grey. Pearson correlation coefficients shown. Example clones shown are 25F_1 clone D6 and 80F_1 clone F7. B) Genome browser view of the chrX:9,779,225-9,957,302 including SHROOM2 gene, an example region that loses methylation in the old clone compared to the young clone. CpG methylation from nanopore sequencing is shown with tracks bounded at 0% and 100% methylation. The first and second tracks show the inactive and active allele in the young clone, respectively. The third and fourth tracks show the inactive and active allele in the old clone, respectively. The fifth track is the phase block, showing the region has been phased into a single block. The sixth track is the CpG island annotation and the seventh track is RefSeq gene annotation. The region around the CpG island of the gene has been expanded showing chrX:9,781,832-9,791,377. D-E) Average methylation over CpG islands of the EMD and ZBTB33 gene across clones from different donors, respectively. Each point is the average methylation over the entire CGI annotation coloured by the donor the clone was derived from. Facets split active and inactive alleles. F) Average methylation over CGIs that do or do not show age-dependent methylation erosion plotted against chronological age of donor. CGIs without methylation erosion were matched in number to those that show methylation erosion, randomly selected 10,000 times and the average of those 10,000 selections is plotted. Pearson correlation coefficients are shown.

To investigate how consistent the age-dependent erosion over CGIs was across different clones, we looked at 3 to 5 independently derived single cell clones from 7 donors (aged 25 to 80 years old). In general, CGIs on the active allele were typically lowly methylated, independent of donor age (Supplementary Figure 2B), whereas CGIs on the inactive allele had higher methylation (Supplementary Figure 2C). Some CGIs, for example the one regulating *EMD*, showed the expected pattern, with low methylation on the active allele in all clones and high methylation on the inactive allele in all clones (Figure 2D). Other genes, for example *ZBTB33*, showed low methylation on the active allele but exhibited loss of methylation in the inactive allele in an age-related manner (Figure 2E). We next compared the methylation over CGIs in clones from the youngest donor (25F) with the equivalent CGIs in clones from all other ages. The active allele was very similar between the clones from the 25 year old and clones from all other ages (Supplementary Figure 2E), whereas the inactive allele showed erosion in many CGIs when compared to clones of other ages (Supplementary Figure 2F). The average methylation across all phaseable CGIs did not significantly decrease with age (Pearson r =-0.34, p-value=0.061, Supplementary Figure 2G). However we were able to identify a subset of 29 CGIs that exhibited a stronger, systematic loss of methylation with age (see methods, Pearson r = −0.78, p-val=1.8×10^-7^, Figure 2F). These CGIs and the genes associated with them will be referred to as age-related escapee genes for the remainder of the manuscript. The methylation did not significantly change with age when randomly sampling an equal number of CGIs on the X chromosome in all of the 10,000 iterations performed (see methods, mean in Figure 2F and null distribution in Supplementary Figure 2H), suggesting that there is a subset of genes on the X chromosome that have an increased propensity to lose methylation with age at their CGIs.

Phasing long read RNA-seq data is non-trivial as most genes do not have informative SNPs in coding regions. Even if there is a SNP present in a coding region, the SNP may be present in exons that are variable between isoforms leading to misleading quantifications of gene expression by looking only at specific isoforms. Due to these technical limitations, we were unable to systematically assess the transcriptional consequences of methylation erosion in the 29 age escapee genes.

### Clustering of age related escapee genes

Given a subset of genes showed substantial erosion of DNA methylation with age, we next asked whether there were hotspots along the chromosome where methylation erosion was more frequently detected. Because not all genes on the X chromosome could be phased, we restricted comparisons to phaseable genes rather than all genes on the X chromosome. The median distance between age-related escapees was 943,288 bp and they were significantly closer to other age escapees than a randomly sampled number of phaseable genes on the X chromosome (see methods, 10,000 iterations with empirical one-sided *p*-value = 0.0037) (Figure 3A,B). We used a common annotation of canonical escapees that were age-independent (Tukiainen et al. 2018) and found that age-related escapees were not significantly closer to canonical escapees than a random subset of phaseable genes (see methods, 10,000 iterations with empirical one-sided *p*-value = 0.75). This suggests that the observed age-dependent methylation erosion is not due solely to the expansion of active regions around canonical escapees on the inactive chromosome (Supplementary Figure 3A,B). However, given that regions with a high density of canonical escapees would contain comparatively fewer methylation differences between the active and inactive allele and therefore would be more difficult to phase using our method, the inability to detect clustering of age-related escapees with canonical escapees may be due to technical limitations.

**Figure 3.**
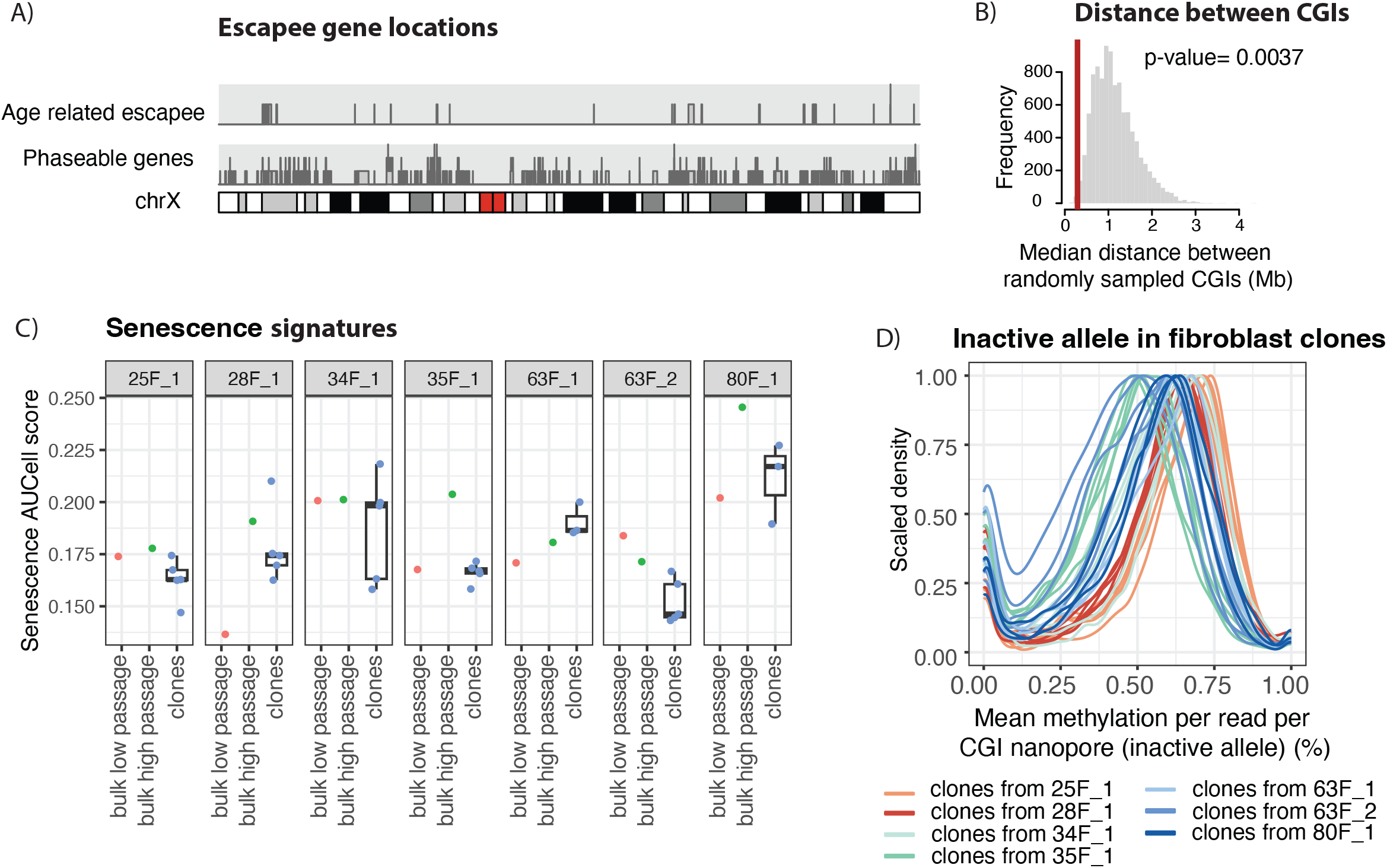
A) Chromosome X karyoplot with locations of all phaseable genes (bottom) and all genes that have been identified as age related escapees (top). B) Null distribution of median distance between randomly sampled phaseable genes on chromosome X, using 10,000 iterations. The red line shows median distance between age-related escapees. Empirical one-sided p-value shown. C) AUCell senescence scores from the top 5% of genes in clones and low and high passage bulk lines (n=56). Multiple clones have been derived from the same donor, so boxplots for clones show median and interquartile range. D) Kernel density estimate of mean methylation over CpG islands calculated per read in reads that have been phased to the inactive allele in clones. Densities are scaled to avoid differences observed due to random variation in total sequencing depth or total number of phaseable regions.

### DNA methylation erosion with age in bulk populations

While it has been reported that methylation phenotype can change during long term culture (Franzen et al. 2021), we observed stable methylation profiles in our fibroblast clones (Supplementary Figure 1B). The cumulative population doublings (CPDs) are similar between young and old donors so it is unlikely that the observed DNA methylation erosion is primarily driven by passage. To assess whether the high number CPDs in the clones induced a senescence phenotype, bulk lines that had not undergone the clonal derivation process were grown in parallel with the clones, providing lower passage (a maximum of 7 passages after receipt from the vendor) and higher passage bulk counterparts (a maximum of 15 passages after receipt from the vendor, similar to clonal lines) (Supplementary table 1). In general, the expression of genes on the X chromosome was very similar between the high and low passage clones (Supplementary Figure 3C). Furthermore, there was not a consistently higher senescence signature in the fibroblast clones compared to lower passage bulk fibroblast cells (linear mixed-effects model with Tukey adjusted contrasts: estimate = –0.001, t = –0.16, p = 0.985) and there was a lower senescence score in clones compared to higher passaged cells (linear mixed-effects model with Tukey adjusted contrasts: estimate = 0.018, t = 2.62, p = 0.033). Together this suggested the observed methylation phenotype was not caused by cells entering senescence (Figure 3C). There was also no significant difference in senescence signatures between higher passage and lower passage fibroblasts further suggesting that senescence signatures were not altered during the timeframe of cell culture used in this study (linear mixed-effects model with Turkey adjusted contrasts: estimate = –0.019, t = –2.19, p = 0.087). Together, this suggests that our observation of age-dependent X chromosome methylation erosion in fibroblast clones is not confounded by an increase in senescence.

We next examined methylation profiles in bulk fibroblast lines that were subjected to limited *in vitro* expansion to determine whether this methylation erosion was observable without the need for clonal derivation. This expanded our sample size beyond the donors we had isolated clones from and resulted in 19 different donors with a more even distribution of ages. Given the variability in the erosion observed from clones from the same donor, we looked at all CGIs and did not just subset to those that had been identified to show age-dependent methylation erosion in our clones. In a young sample that has been sequenced deep enough, the expectation would be that the average methylation of any given CGI should be around 50%; if the inactive allele is fully methylated and the active allele is fully unmethylated. When erosion occurs in older samples, there should be an observable shift away from 50%. However in practice, the inactive allele is not 100% methylated at every CGI meaning that average methylation over both alleles is not 50%. Furthermore, it is apparent from the clonal data that different cells are heterogeneous both in the amount of methylation erosion and which CGIs show erosion. Together, this means the average methylation over CGIs in bulk does not have the resolution to identify methylation erosion with age (Supplementary Figure 4A,B).

However, nanopore reads are long enough to span an entire CGI. Each read that spans a CGI will come from either the inactive or the active allele in a specific cell and provides a measure of the methylation over that CGI. This allows read based quantification of methylation, where each read acts as a proxy for either the active or the inactive allele in an individual cell. CGIs on reads from the active allele will be very lowly methylated showing an average methylation near 0%. Conversely, CGIs on reads from the inactive allele will have much higher methylation with an average methylation closer to 100%. Given that we did not see many age-related changes in methylation on the active allele in the fibroblast clones, we expect many reads with low methylation irrespective of age. However, if the erosion is detectable using this read-based method, we should see a downward shift in the average methylation per read of those reads with high methylation. First we looked in only the inactive reads from our phased clonal data, looking at all reads and not just ones that overlapped genes that have been identified as age related escapees in our clonal lines. The mean methylation decreases across the population of inactive reads with age (mean read based methylation per donor with age, Pearson r = –0.37, p-value = 0.037, Figure 3D). There was also an observable peak around 0%, likely representing canonical age-independent escapees or genes that have no methylation on the inactive allele. We next turned to the unphased clonal data to see if we were able to identify these erosion patterns using this read based method without separating out the active and inactive alleles. When looking across all CGIs on the X chromosome, we saw the expected peak of reads with 0% methylation that are derived primarily from the active allele. Notably, we also observed the age-related downward shift of the CGIs with higher methylation, consistent with age-associated erosion (mean read based methylation per donor with age: Pearson r = –0.49, p-value = 5.5×10^-3^, Supplementary Figure 4C).

Given our ability to observe these dynamics in the clonal data, we expanded our sample size beyond the donors we had isolated clones from to the bulk lines with a more even distribution of ages. Again looking across all CGIs and not just the ones that exhibited age-dependent methylation erosion in the clones, we observed an age-dependent reduction in methylation levels in the fibroblast bulk lines (Figure 4A). To identify which parts of the methylation distribution changed with age, the average methylation per CGI per read was ordered from least to most methylated for each sample. The methylation values at each decile were extracted from this ordered list to create a 10 number profile per sample that summarised the methylation distribution for each sample. From this, we identified that deciles 5, 6, 7 and 8 showed significant loss of methylation in an age-dependent manner indicating that the greatest age-related change in methylation was seen for CGIs that had an average methylation of 35% to 64% (Figure 4B, Pearson r −0.48, −0.54, −0.54, −0.54, and Bonferroni adjusted p-value =3.8×10^-2^, 1.9×10^-2^, 1.9×10^-2^, 1.9×10^-2^ respectively). This clearly shows that CGI methylation erosion on the inactive X chromosome can be observed without the extensive passaging that occurs during clonal expansion. Together, this suggests that our observation of age-dependent X chromosome methylation erosion over CGIs in fibroblasts is not confounded by long term passage and is representative of differences in the donor cells that were present before clonal expansion.

**Figure 4.**
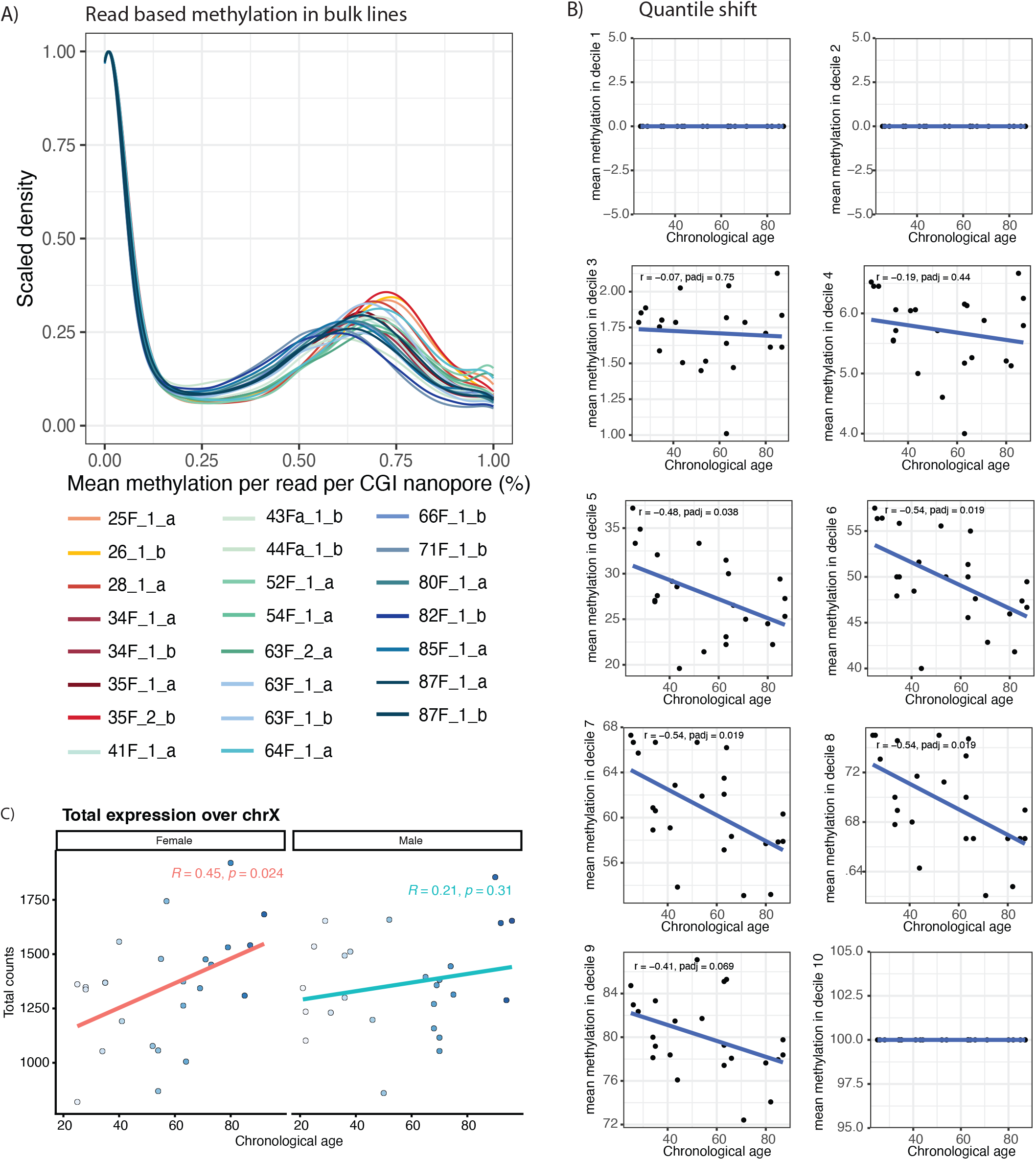
A) Kernel density estimate of mean methylation over CpG islands calculated per read in unphased non-clonal lines, showing methylation patterns of both the active and the inactive alleles (a and b denote culture method). Densities are scaled to avoid differences observed due to random variation in total sequencing depth. B) Mean methylation of CGI per read per donor split into deciles. For each sample the mean methylation of reads that fall into the decile is plotted. Pearson correlation coefficient and p-value shown in each panel. Methylation at deciles 5 to 8 decreases with age. C) Total counts of expressed age-associated escapee genes (n=22) split by sex. Counts normalized using Trimmed Mean of M-values. Pearson correlation coefficients and p-values shown.

Given that the repression of the inactive X chromosome is also dependent on other epigenetic modifications and not just DNA methylation (Galupa and Heard 2018), this observed loss of methylation at CGI does not necessarily mean that there will be transcriptional reactivation. We therefore sought to query whether any of these genes show evidence of disrupted transcription in bulk transcriptomic data from dermal fibroblasts across a range of ages. Since we cannot effectively separate active and inactive transcripts, we looked at expression in males and females under the assumption that the patterns observed in males were representative of the active allele and these patterns were similar in males and females. Then, any observed differences in expression between males and females would result from differences arising on the inactive allele with age. Given the heterogeneity we observed in age-related erosion of CGI methylation both within donors and between donors in our clonal data, a single gene may not change in every donor. Therefore, we looked at total transcription across all genes to identify increased signal in aggregate over these genes. There was no relationship between total gene expression from the X chromosome and age (female Pearson r = 0.063, p-value = 0.76, male Pearson r = 0.042, p-value = 0.71, Supplementary Figure 4D). We next looked if there was observable signal specifically in the genes identified as age-related escapee genes in our clonal data. Some genes do not appear to be expressed from either allele, perhaps suggesting the absence of necessary transcription factors in this cell type. This left 16 genes from our list (above mean cpm 0, Supplementary figure 4F). With the remaining age-related escapee genes, there was an increase in total expression with age in females (Pearson r = 0.45, p-value = 0.024, Figure 4C) but not in males (Pearson r = 0.21, p-value = 0.31, Figure 4C). This suggested that for some of these escapee genes there was transcriptional upregulation. This is consistent with previous work in mice (Hoelzl et al. 2025; Gadek et al. 2025) and indicated that if the correct combination of chromatin context and transcription factors are present, increase of X dosage is possible.

## Discussion

The X chromosome has presented a challenge for genomic analyses due to the differences between the active and inactive alleles and the mosaicism of X inactivation in human tissues. While short read technologies provide a fragmented glimpse of the underlying biology, the advent of long-read sequencing technologies has fundamentally altered this landscape. They provide multi-kilobase reads and enable the effective phasing of maternal and paternal alleles, while also providing the ability to detect native DNA modifications for high-resolution epigenetic profiling. This synergy allows for the discovery of new insights into X chromosome biology that were previously not possible (Vollger et al. 2025; Dubocanin et al. 2025; Spix et al. 2025).

Here we used long-read sequencing of human dermal fibroblasts to study changes on the X chromosomes with age. Given that tissues are heterogeneous with respect to which parental allele is inactivated, we used clonally derived cell lines to identify regions that systematically show age-dependent alterations in methylation. On average 40% of the X chromosome could be confidently assigned to the active or inactive allele. Since our assignment strategy relies heavily on CGI and gene-body methylation, this assignable fraction was enriched for genes and CpG islands relative to the rest of the chromosome. When comparing the active and inactive alleles, we observed the expected differences in methylation between CGIs and gene bodies and, for genes which contain heterozygous SNPs, were able to observe differences in expression. We used these features to assign haplotypes within phase blocks to either the active or the inactive allele. We then interrogated how the methylation patterns of the inactive X chromosome changed with age. Some CGIs showed erosion in only a single clone or in a donor specific manner, while others systematically lost methylation in multiple clones and donors in an age-dependent manner. The CGIs that showed methylation erosion were clustered genomically suggesting that there may be larger scale chromatin conformation differences that may give these regions an increased propensity to change. Importantly, the CGI erosion did not affect the active vs inactive allele assignment as the assigned portions of the X chromosome did not correlate with the age of the donor. The initial observation of age-dependent methylation erosion was made in the clonal cohort which was composed of 7 donors and 31 clones. This erosion was recapitulated in 19 bulk, non-clonally derived donors, suggesting the phenotype is not solely a consequence of long term *in vitro* cell culture.

The observed erosion may be contributing to age-related changes in cell function that differ between older men and women. This may have a positive or negative impact depending on the selection pressures, tissue of origin and context. In mice for example, Hoelzl *et al* find age-related transcriptional escape in various organs, including the heart, lung, liver, brain, spleen, kidney, and muscle, and observe that age-escapees demonstrate increased chromatin accessibility and tend to cluster at distal chromosomal regions (Hoelzl et al. 2025). In particular, they observe escape of *Ace2* in the lung of mice and *Ace2* overexpression has been shown to be protective against pulmonary fibrosis in rats (Shenoy et al. 2010). Gadek *et al* find age-related escape from inactivation of the X chromosome gene *Plp1* in females. When *Plp1* was overexpressed in the dentate gyrus, they observed improved cognition irrespective of sex (Gadek et al. 2025). Some neurodegenerative diseases also have a sex bias and occur more in older individuals. In those that are more prevalent in men than women and have increased incidence with age, such as Parkinson’s disease (Moisan et al. 2016), these overexpressed genes may be protective. However, overexpression due to X chromosome inactivation erosion may also be detrimental. In diseases that are more prevalent in women than men and have increased incidence with age, such as Alzheimer’s disease (Beam et al. 2018), these overexpressed genes may be exacerbating incidence. Furthermore the X chromosome contains the largest number of immune related genes in the human genome (Bianchi et al. 2012) with X-linked genes that are known drivers of autoimmune disease (G. J. Brown et al. 2022), and indeed some autoimmune diseases are far more common in women than in men with age (Conrad et al. 2023)

We have shown that X chromosome methylation erosion is associated with an upregulation of gene expression when looking at total transcription of the age-dependent escapee genes in bulk data. We hypothesise that this erosion of X chromosome inactivation contributes to the differences in diseases which have different incidences between the sexes with age. In some cases, single genes may be driving these effects (Gadek et al. 2025; Hoelzl et al. 2025). However in other cases, the small upregulation of multiple genes on the otherwise inactive X chromosome may collectively lead to increased disease susceptibility. This is consistent with a recent human allele-specific expression study, which found that age-associated escape from XCI is typically modest in magnitude across many genes rather than major derepression at a few loci (Rocca et al. 2026). Traditional methods to identify disease driving genes often rely on differential gene expression analysis between disease and control samples or genome wide association studies. These methods are very effective at identifying single gene drivers that contribute to the majority of the phenotype. However they would be less effective at identifying genes that become upregulated due to large-scale chromatin dysregulation, such as the X chromosome erosion described here. Therefore the methodology described in this paper could allow new disease influencing genes to be discovered.

## Materials and Methods

### Cell culture fibroblasts

Primary human dermal fibroblasts (HDFs) were obtained from three commercial sources: the Coriell Institute for Medical Research (Camden, NJ, USA; donor/catalog IDs GM00038, AG03198, AG06299, AG08468, AG11362, and GM03525), PromoCell (Heidelberg, Germany; donor/catalog IDs 486Z014.1 and 482Z012.3), and Cell Applications, Inc. (San Diego, CA, USA; donor/catalog ID 2049). Donors ranged in age from 25 to 80 years at the time of biopsy (Supplementary Table 1). Two culture methods were used. The following cell lines were cultured using cell culture method a: 25F_1, 28F_1, 34F_1, 35F_1, 41F_1, 52F_1, 54F_1, 63F_1, 63F_2, 64F_1, 80F_1, 85F_1 and 87F_1. For culture method a, cells were cultured in DMEM/F-12 with GlutaMAX (Gibco, 31331-028) supplemented with 10% fetal bovine serum (Thermo Fisher, A5256801), 50 µM µM β-mercaptoethanol (Gibco, 31350-010), [1×] MEM non-essential amino acids (Gibco, 11140-050), and 16 ng/mL fibroblast growth factor 2 (FGF2) (Qkine, Qk053), on tissue-culture-treated plastic plates and flasks. Cells were maintained at 37°C in a humidified incubator with 5% CO,. The following cell lines were cultured using cell culture method b: 26F_1, 35F_2, 34F_1, 43F_1 44F_1, 63F_1, 66F_1, 71F_1, 82F_1 and 87F_1 For culture method 2, cells were cultured in complete growth medium consisting of DMEM basal medium (Gibco, 11965-092) supplemented with 15% fetal bovine serum (Omega Scientific, FB-01), 1% non-essential amino acids (Gibco, 11140-050), and 1% GlutaMAX (Gibco, 3505-061). Cells were maintained at 37°C in a humidified incubator with 5% CO,. The different culture methods did not affect CGI methylation during the culture period used in this study (Supplementary Figure 4E). Cells were passaged twice weekly using TrypLE Express (5 min, 37 °C) and cryopreserved in CryoStor cryopreservation medium (Sigma Aldrich, C2874). Low-passage lines were collected within a few passages of the initial expansion from the vendor-supplied cryovial.

### Clonal derivation fibroblasts

For clonal derivation, cells were dissociated with TrypLE and single cells were isolated by fluorescence-activated cell sorting directly into individual wells of 96-well plates containing culture medium supplemented with [1×] penicillin-streptomycin (kept only for a week to avoid contaminations derived from sorting). Clones were cultured in DMEM/F-12 with GlutaMAX (Gibco, 31331-028) supplemented with 10% fetal bovine serum (Thermo Fisher, A5256801), 50 µM µM β-mercaptoethanol (Gibco, 31350-010), [1×] MEM non-essential amino acids (Gibco, 11140-050), and 16 ng/mL fibroblast growth factor 2 (FGF2) (Qkine, Qk053), on tissue-culture-treated plastic plates and flasks. Clones were maintained at 37°C in a humidified incubator with 5% CO,. Single-cell-derived colonies were allowed to recover for 1–3 weeks until they covered the majority of the well surface area. Wells containing viable clonal outgrowths were identified by visual inspection, dissociated with TrypLE, and sequentially expanded through 12-well, 6-well plates, 10 cm and 15 cm dishes, depending on the scale of expansion required and individual clone growth rate, to generate sufficient cell numbers for high-molecular-weight DNA extraction for nanopore sequencing. Passage numbers, culture dates, and cell yields for all clonal and bulk lines are provided in Supplementary Table 1.

### DNA Sequencing

Genomic DNA was isolated using either the Monarch Spin gDNA Extraction Kit (New England Biolabs) with Proteinase K and RNase A treatment or the Quick-DNA/RNA Microprep Plus Kit (Zymo Research). DNA concentration was quantified using Qubit, purity assessed by NanoDrop spectrophotometry and DNA integrity was determined using an Agilent TapeStation.

Libraries were prepared without fragmentation. Genomic DNA was either prepared for adapter ligation in a single end-repair and A-tailing reaction (NEBNext® Companion Module v2 for Oxford Nanopore Technologies® Ligation Sequencing) or sequentially as end-repair (NEB FFPE DNA Repair Mix in FFPE v2 buffer), 1X SPRIselect (Beckman), followed by A-tailing (Watchmaker Genomics DNA Library Prep kit). Libraries were prepared using ligation sequencing kit V14 SQK-LSK114 (Oxford Nanopore Technologies) and loaded onto one R10.4.1 PromethION FLO-PRO114M or FLO-PRO114M-HD flow cell per sample. Where required due to pore blocking, flow cells were flushed using wash kit EXP-WSH004 and reloaded after approximately 24 and 48 hours of sequencing.

### RNA Sequencing

For long-read sequencing, total RNA was isolated using the Quick-DNA/RNA Microprep Plus Kit (Zymo Research) with on-column DNase I digestion, following the manufacturer’s instructions. RNA concentration was quantified using Qubit, purity assessed by NanoDrop spectrophotometry and RNA integrity was determined using an Agilent TapeStation.

Libraries were prepared using the Oxford Nanopore Technologies (ONT) cDNA-PCR Barcoding Kit V14 (SQK-PCB114.24). Reverse transcription was performed with Maxima H Minus Reverse Transcriptase for 90 min at 42 °C, a modification of the standard protocol. cDNA amplification was performed in a total volume of 25 µl containing 12.5 µL of 2 x Ex Premier DNA Polymerase (TaKaRa), 2.5 µl of each barcoding primer (BP01-24, 2 µM) and 10 µl cDNA. Cycling conditions consisted of an initial denaturation at 94 °C for 1 min; 14 cycles of 98 °C for 10 s, 68 °C for 6 min; followed by a final extension at 68 °C for 6 min. After excess primer removal with Thermolabile Exonuclease I (NEB), samples were purified with 0.68X AMPure XP SPRI beads. Samples were grouped into three pools prior to an additional 0.68X Ampure XP SPRI size selection. Rapid sequencing adapters were added to each pool and loaded onto three R10.4.1 PromethION FLO-PRO114M flow cells per pool.

For Illumina short-read RNA sequencing, the Quick-DNA/RNA Microprep Plus Kit (Zymo Research) was used to isolate total RNA following manufacturer guidelines. The Illumina NovaSeq platform was used to generate 150 bp paired-end whole genome sequencing with 40-76 million reads per library.

### Genomic alignment and variant calling

GRCh38 was used as the reference genome. For nanopore whole genome sequencing, reads were trimmed using PyChopper (v2.7.10), basecalling was performed using dorado (v1.0.0) with base calling model dna_r10.4.1_e8.2_400bps_sup@v5.2.0. Reads were aligned to the reference genome using minimap2 (v2.30). Modkit (v0.6.1) was used to extract methylation information from bam files and perform downstream processing.

Single nucleotide variants in the clones were called using Clair3 (v1.0.5). Variants overlapping simple repeats and with base quality score less than 20 were removed. True germline variants were expected to be observed at a heterozygous variant allele fraction, so a one-sided exact binomial test was used on the number of variant supporting reads and total number of reads across samples to ensure only true germline variants were kept for phasing, as described previously (Coorens et al. 2021).

### Transcriptomic alignment and gene quantification

GRCh38 was used as the reference genome. For nanopore whole genome sequencing, reads were trimmed using PyChopper (v2.7.10), basecalling was performed using dorado (v1.0.). Reads were aligned to the reference genome using minimap2 (v2.30). When genes contained SNPs, reads were phased using phase block assignment from the DNA sequencing. Gene level counts were quantified using Isoquant (v3.7.0) (Prjibelski et al. 2023) using Ensembl annotation v108. Counts were normalised using Trimmed Mean of M-values.

### Phasing method

To phase using genomic variants, only reads with a mapping quality greater than 60 were used. Since nanopore sequencing data has a high rate of indel occurrence, the number of sequence mismatches and indels as a proportion of the total read length was calculated. If this exceeded 3%, these reads were not used for downstream phasing but were re-encorporated once phase blocks were determined. Whatshap (Patterson et al. 2015) was used to phase genomic variants using SNVs and to tag reads based on this phasing.

Once genomic phase blocks were established, methphaser (Fu et al. 2024) was used to extend phase blocks using distinct methylation patterns between the two alleles and further tag the reads. Again, only reads containing indels and SNPs that accounted for less than 3% of the total read length were used. In some cases there was insufficient evidence to extend the phase blocks despite there being reads that span two adjacent blocks, for example if local read depth is very low. In these cases, the reads needed to be removed to prevent these reads being assigned to both haplotypes.

These phase blocks then needed to be assigned to the active or inactive chromosome. To do this, the average methylation for each phase block is determined over CpG islands and over the gene body. Publicly available CpG islands annotations were used (Illingworth et al. 2010). This annotation did not contain a CpG island for *Xist*. Given the importance of Xist in X inactivation, a region centered on the transcriptional start site was manually annotated (X:73852864-73850559).

To validate this as a region that is informative for phasing, the raw methylation patterns were inspected. It contained one highly methylated allele and one unmethylated allele. While it is possible to do this for all genes without a CpG island, inappropriately specifying regions can lead to switch errors in phasing so *Xist* was the only gene that was manually annotated.

The following rules were used for assignment in a sequential manner:

1. If any CpG island within the phase block had a methylation difference greater than 25%, then the block with higher methylation was assigned to the inactive allele and the block with the lower methylation was assigned to the active allele. All the CpG islands within the phase block must agree in direction for the blocks to be assigned. If some CGIs had higher methylation and others had lower methylation on the same allele, this phase block was not assigned.
2. Some blocks contained gene bodies where the CpG island of the gene was contained in another block that was assigned by rule 1. In these cases, the gene body methylation in the unassigned block was considered. If the gene body methylation difference was greater than 25% between alleles, then the gene body with higher methylation was assigned to the active allele and the gene body with lower methylation was assigned to the inactive allele.
3. Calculate the allelic ratio of all genes within the phase block that had a minimum total expression of 10 reads irrespective of allele. Genes within the phase block that had an allelic ratio around 0.5 were considered to be canonical escapees and were not used to influence assignment. If the allelic ratio for all assigned genes within a phase block was greater than 0.8 or less than 0.2, then the allele with the higher allelic ratio was considered to be the active allele.

### Identification of age related escapee genes

The average methylation over all phaseable CGIs on the X chromosome was calculated per clone. Linear modelling using the R package *limma* (Ritchie et al. 2015) was used to assess how each CpG island changed with respect to covariate of age and the commercial vendor the cell line was purchased from. Differentially methylated CpG islands were identified, with statistical significance threshold set as a Benjamini-Hochberg (BH) adjusted p-value < 0.05. Age-related escapees must significantly change with age and have a negative age coefficient less than −0.1. They must not be significantly changed between different commercial vendors.

### Random sampling of methylation at CGIs

To identify whether the age-related erosion of CGI was greater than expected by change, a null distribution of an equal number of randomly sampled phaseable genes (29) was generated. This random sampling was performed 10,000 times only using genes that did not show evidence of escape. For each iteration, the median methylation was calculated across those genes per sample and a Pearson correlation coefficient was calculated against the age of the donor the clone was derived from. The null distribution was formed from these 10,000 correlations coefficients. The difference between the null distribution and the correlation of age escapees was quantified using an empirical two-sided p-value using (b + 1) / (m + 1), where b is the number of correlation coefficients from the randomly sampled CGIs that have an absolute correlation greater than or equal to the correlation of the age-related escapees and m is the number of iterations (Phipson and Smyth 2010).

### Clustering analysis

To identify whether escapee genes were genomically clustered, the nearest gene for every escapee was identified with the distance defined using the start position of both genes. The median distance for all 29 escapees was then calculated. The null distribution was generated using 10,000 randomly sampled phaseable genes. For each iteration, the distance to the nearest gene was calculated for each of the random genes and the median distance of all genes in that random set was calculated. To test whether the age-related escapes were closer to other age related escapees, an empirical two-sided p-value was calculated using (b + 1) / (m + 1), where b is the number of median distances from the randomly sampled CGIs that have an absolute median distance greater than or equal to the median distance of the age-related escapees and m is the number of iterations (Phipson and Smyth 2010). The same method was used to test whether the age-related escapees were closer to canonical escapees (as annotated by (Tukiainen et al. 2017)) than randomly sampled phaseable genes.

### Senescence signatures

Senescent cell signatures in the clones and bulk lines were calculated using the SenMayo gene signature (Saul et al. 2022). Per-sample scores were calculated using AUCell (v1.32.0) on raw counts using the top 5% rank threshold. Ties between gene ranks were resolved randomly. This generated the senescence score. Senescence scores were modelled by linear mixed-effects regression with sample type (clone or bulk) modelled as a fixed effect and donor modelled as a random effect. Estimated marginal means and all pairwise contrasts between sample types were calculated with emmeans (v2.0.3), with p-values adjusted for multiple comparisons by Tukey’s method.

## Author contributions

JI, IK, FS and WR designed the study. Cell lines used in this study were coordinated and managed by JT and IK. IK and JC cultured cells for clonal data and RC performed single cell sorting prior to clonal expansion. IK, AP, AL, and JST cultured cells for bulk data. IK performed nucleic acid extractions for all data. NP and MFA performed library preparation and sequencing and PC coordinated sequencing submission. FK performed core bioinformatic processing of raw data. JI and IK conceived and implemented phasing methodology. JI performed downstream data analysis, with input from IK. RA and SAM contributed to general scientific discussion. The manuscript was written by JI, IK, FS and WR with input from all authors.

## Acknowledgments

We thank all members of the Reik lab as well as Ken Raj and Fynn Krause for their scientific input and helpful discussions. We thank Chris Hammer, Sam Drake, Laura Martin, Amy Peterson, Ana Granat for sharing the bulk nanopore sequencing method that was used. We thank Alice Santambrogio for technical support and assistance with cell culture. We thank Manuel Serrano for his review of the manuscript.

---

**Supplementary Figure 1.**
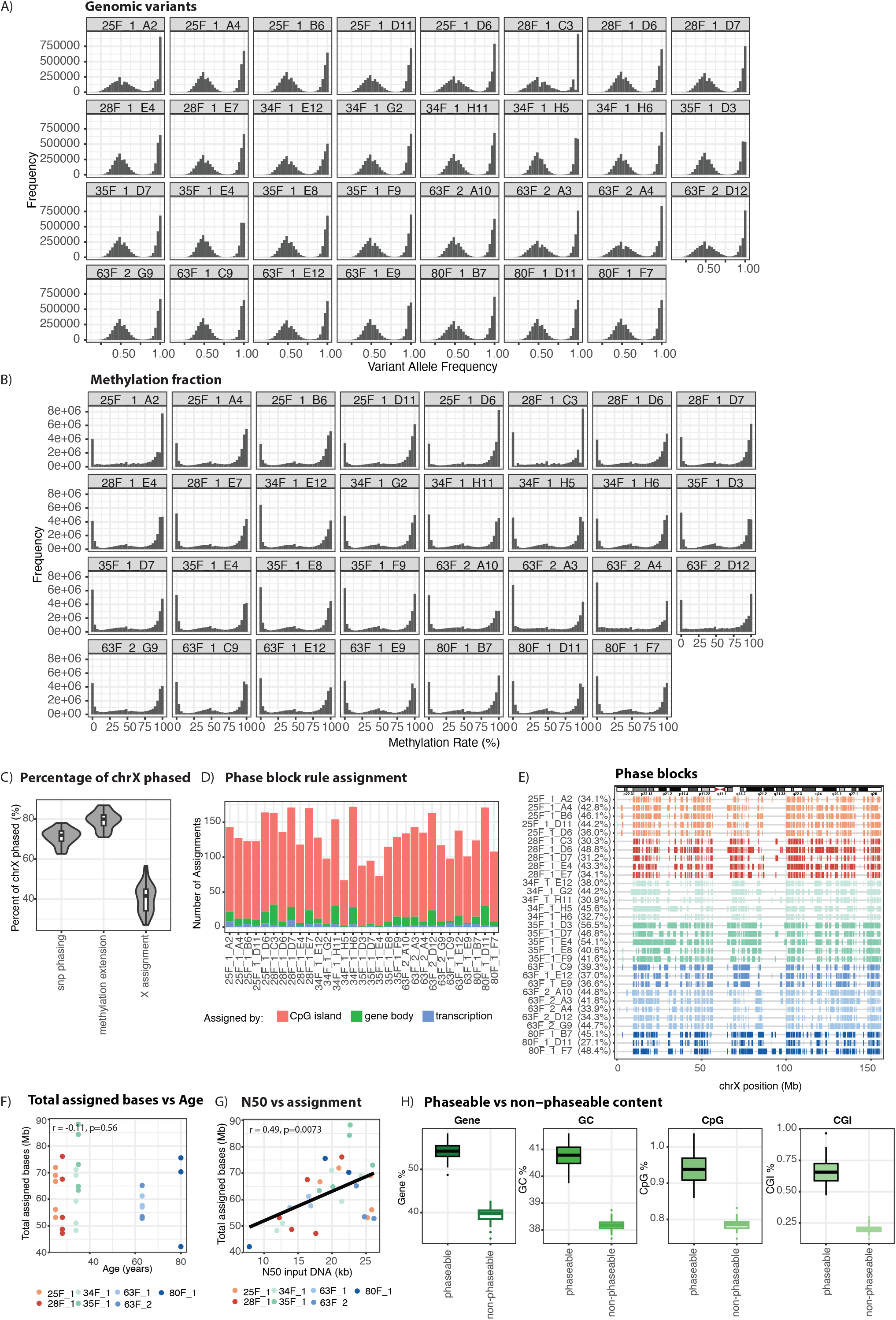
A) Variant allele frequency (VAF) histogram of variants faceted by clones. B) Histogram of methylation (methylated C’s/total C’s) at all CpGs across all clones. C) Violin showing distribution of the proportion of chromosome X phased in each step of the haplotype phasing method across all 31 clones. D) Number of phase blocks assigned by CpG island methylation, methylation differences across gene bodies spanning phase blocks and transcription. E) Phase block locations across all the clones. F) Total number of assigned bases compared to age of the donor the clones were derived from. Pearson correlation coefficient and p-value shown. G) Total number of assigned bases compared to the length of the shortest read where all reads of that length or longer account for at least 50% of the total base pairs sequenced (N50). Pearson correlation coefficient and p-value shown. H) Gene, GC, CpG and CpG island in phaseable vs non-phaseable regions of the X chromosome.

**Supplementary Figure 2.**
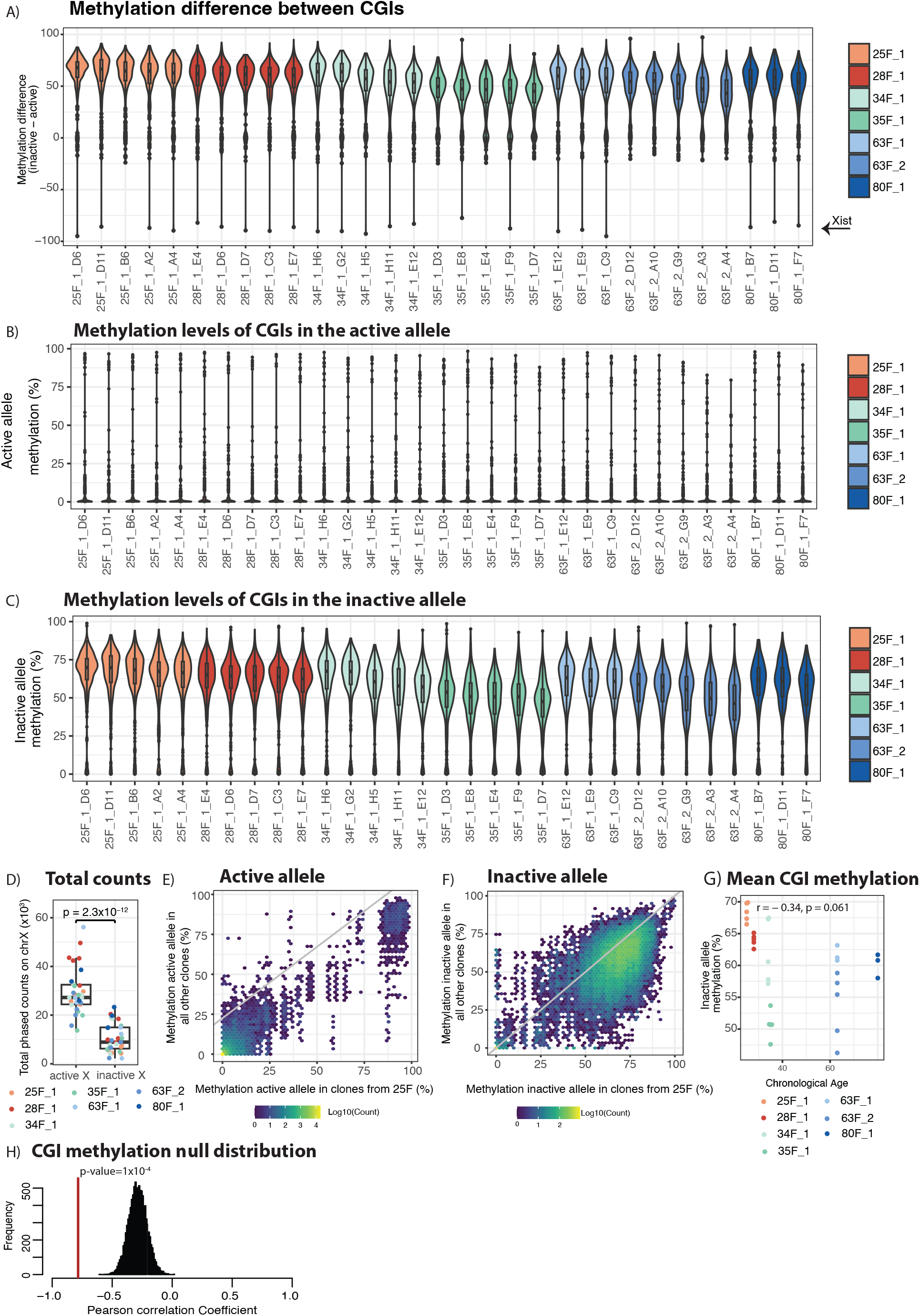
A) Violin showing distribution of differences in average methylation over phaseable CpG islands between the inactive and active allele within each clone. Violins scaled to equal width. Overlaid boxplots show median and interquartile range. B-C) Violin showing distribution of average methylation over phaseable CpG islands on the active and inactive allele, respectively. Violins scaled to equal width. Overlaid boxplots show median and interquartile range. D) Total counts from phased reads on the active and inactive allele. Counts normalised based on total counts from the entire genome. P-value from paired t-test shown. E-F) Mean methylation over all phaseable CpG islands on the active allele and the inactive allele, respectively, in all clones derived from 25F compared to clones from all other donors. E) Mean methylation over all CGIs against chorological age of donor, irrespective of whether they show consistent age-dependent loss of methylation. Pearson correlation coefficient and p-value shown. F) Histogram of Pearson correlation coefficients of median methylation of randomly sampled CGIs and age which form the null distribution. The red line shows the correlation coefficient for the median methylation of age-related escapee CGIs with age. Empirical one-sided p-value shown.

**Supplementary Figure 3.**
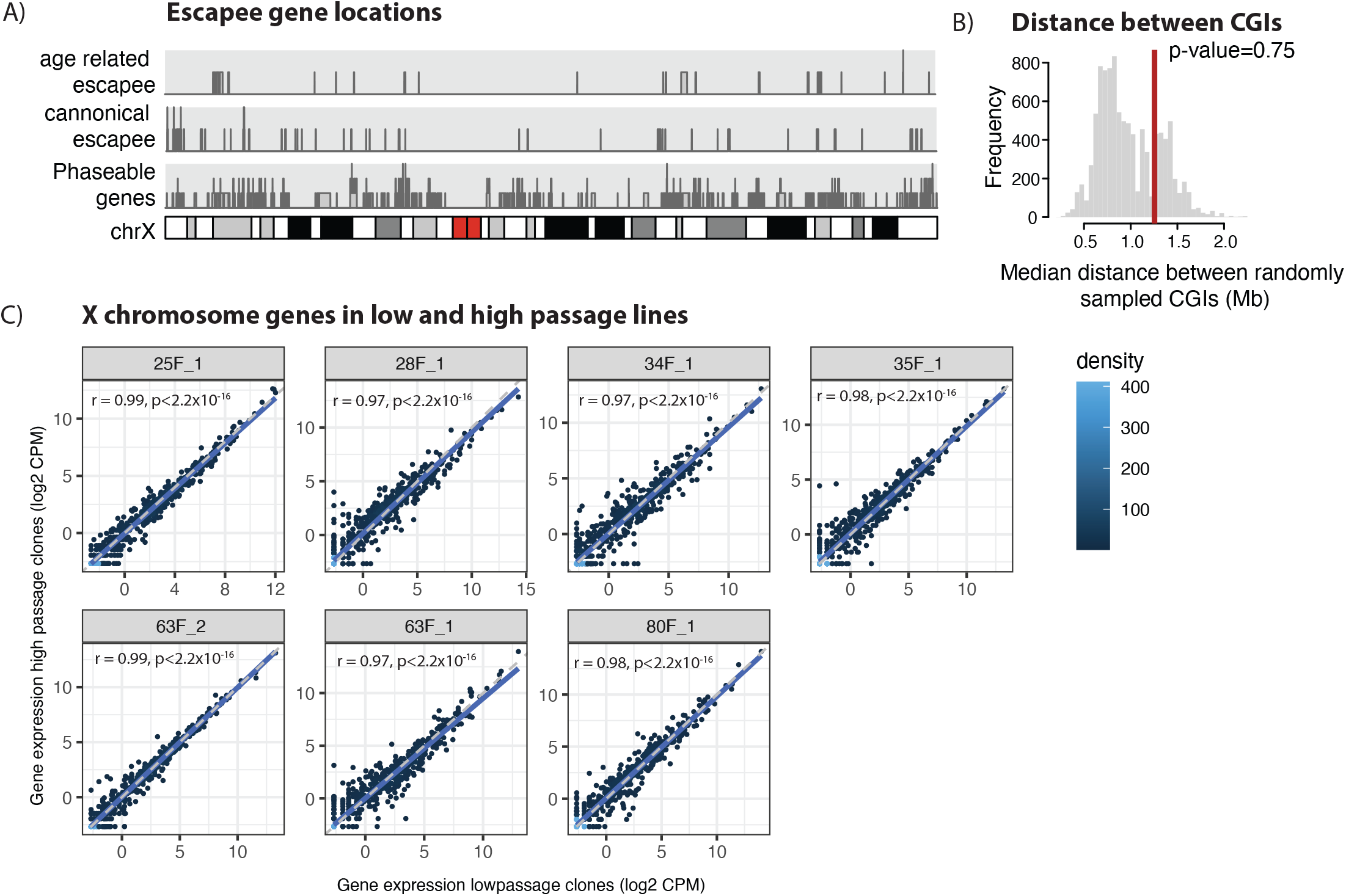
A) Chromosome X karyoplot with locations of all phaseable genes (bottom), canonical escapee annotated genes (middle) and all genes that have been identified as age related escapees (top). B) Null distribution of median distance between randomly sampled phaseable genes on chromosome X and canonical escapee genes, using 10,000 iterations. The red line shows median distance between canonical escapees and age related escapees. Empirical one-sided p-value shown. C) Gene expression of genes on the X chromosome between low and high passage bulk cell lines. Pearson correlation coefficient shown.

**Supplementary Figure 4.**
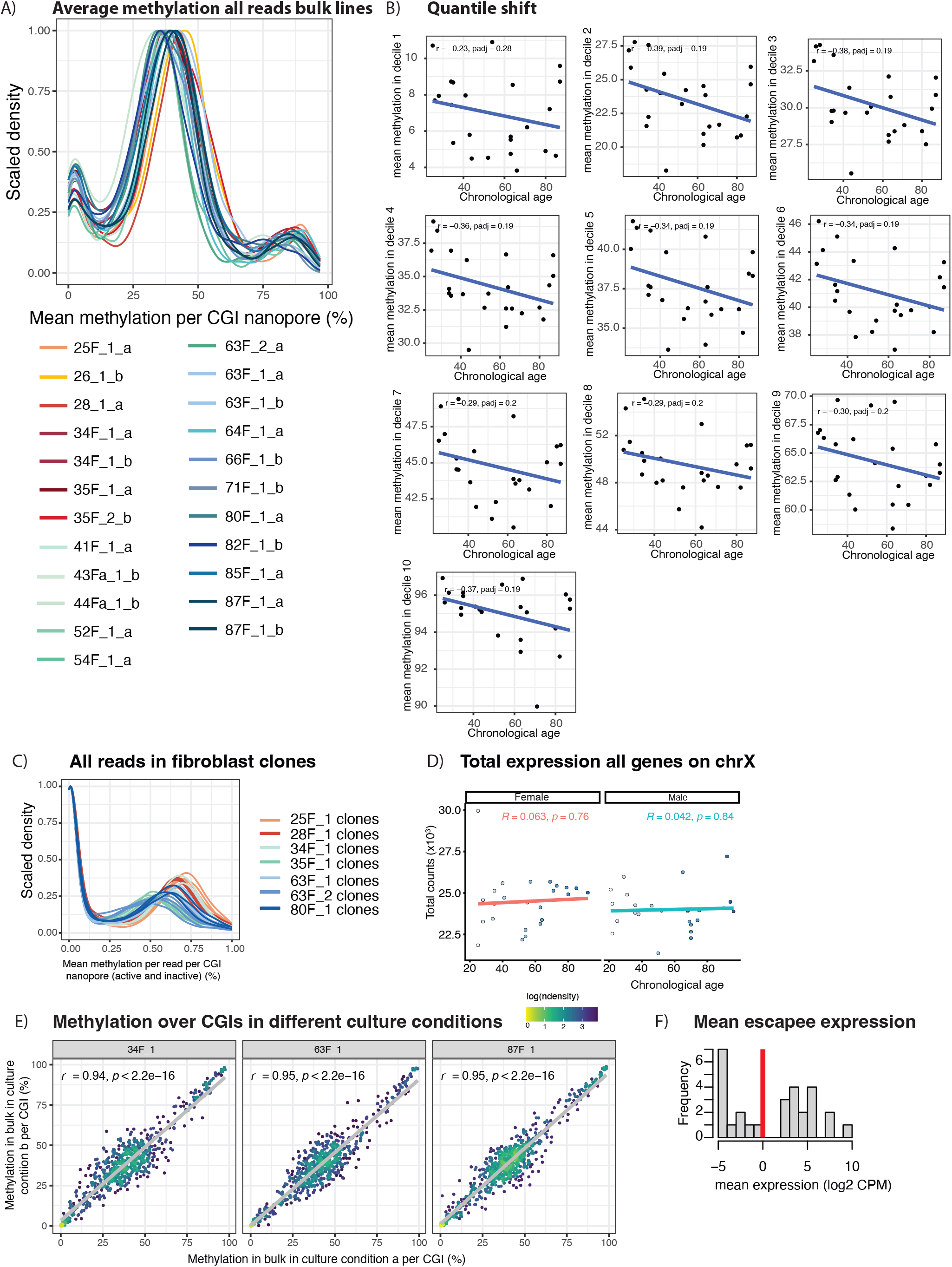
A) Kernel density estimate of mean methylation over CpG islands averaged over all reads in bulk unphased data (a and b denote culture method). Densities are scaled to avoid differences observed due to random variation in total sequencing depth. B) Mean methylation of CGIs averaged over all reads per donor in bulk data split into deciles. For each sample the mean methylation of CGIs that fall into the decile is plotted. Pearson correlation coefficient and Bonferroni corrected p-value shown in each panel. C) Kernel density estimate of mean methylation over CpG islands calculated per read in unphased clones. Densities are scaled to avoid differences observed due to random variation in total sequencing depth. D) Total counts of all genes on the X chromosome split by sex. Counts normalized using Trimmed Mean of M-values. Pearson correlation coefficient and Bonferroni corrected p-value shown. E) Mean methylation over CGIs on the X chromosome in cell lines culture in two different media conditions. Points are coloured by local log-scaled normalised point density. Pearson correlation coefficients shown. F) Mean log2 CPM of gene expression of escapee gene. Red line indicates the expression threshold above which the mean gene expression needed to be considered expressed.

## Notes

### Competing Interest Statement

All of the authors are employees of Altos Labs.

### Summary of Updates

The figures in the initial upload did not leave 1 inch at the top of the page to allow for the licensing statement. This has now been amended to leave 1inch at the top of the page so that the license information should not overlay the figures and the figures are unobscured. Also figure 3 had subpanels A,B,E,F which is changed to A,B,C,D

### Funding

- Altos Labs

